# Characterization of mitochondrial health from human peripheral blood mononuclear cells to cerebral organoids derived from induced pluripotent stem cells

**DOI:** 10.1101/2020.06.05.135392

**Authors:** Angela Duong, Alesya Evstratova, Adam Sivitilli, J. Javier Hernandez, Jessica Gosio, Azizia Wahedi, Neal Sondheimer, Jeff L. Wrana, Jean-Martin Beaulieu, Liliana Attisano, Ana C. Andreazza

## Abstract

Mitochondrial health plays a crucial role in human brain development and diseases. However, the evaluation of mitochondrial health in the brain is not incorporated into clinical practice due to ethical and logistical concerns. As a result, the development of targeted mitochondrial therapeutics remains a significant challenge due to the lack of appropriate patient-derived brain tissues. To address these unmet needs, we developed cerebral organoids (COs) from induced pluripotent stem cells (iPSCs) derived from human peripheral blood mononuclear cells (PBMCs) and monitored mitochondrial health from the primary, reprogrammed and differentiated stages. Our results show preserved mitochondrial genetics, function and treatment responses across PBMCs to iPSCs to COs, and measurable neuronal activity in the COs. We expect our approach will serve as a model for more widespread evaluation of mitochondrial health relevant to a wide range of human diseases using readily accessible patient peripheral (PBMCs) and stem-cell derived brain tissue samples.

## INTRODUCTION

Mitochondrial dysfunction plays a crucial role in a wide range of human diseases^1,2^. The impact of aberrant mitochondrial activity is vast and disease burden will continue to increase unless we uncover their etiology and identify precise mitochondrial targets appropriate for the development of effective treatments. The brain consumes 20% of the total energy budget to power neuronal activity^3-5^. As a result, chronic mitochondrial dysfunction can have profound effects on neurotransmission and contributes to unwanted changes in neuronal circuits that underlie cognition, memory and other forms of neuronal plasticity^3,6,7^. The translation of this knowledge towards the development of effective drugs that target brain mitochondrial dysfunction remains at an early stage. This stagnation reflects a lack of adequate, functional, patient-derived models to study mitochondrial health and neuronal activity simultaneously. To date, tools for studying mitochondrial dysfunction have largely relied on postmortem brain samples, animal models or two-dimensional neuronal systems^8,9^. While these tools have been beneficial, they do not translate well into clinical applications, largely due to the lack of complex functions and neural circuits. These limitations restrict the accurate prediction of patient responses and screening of mitochondrial therapeutic compounds. In the absence of properly developed, patient-derived brain models capable of interrogating mitochondrial health, substantial barriers in testing etiological hypotheses and developing targeted mitochondrial therapeutics will continue to persist.

Cerebral organoids (CO) have become an essential tool for evaluating human brain development and diseases^10^. The ability of CO to differentiate into many cell types and self-organize three-dimensionally makes them a unique and powerful tool for disease modelling and evaluation of mitochondrial health and neuronal activity^10,11^. COs have already been developed and extensively characterized using induced pluripotent stem cells (iPSCs) derived from human dermal fibroblasts^11–13^. However, this approach has drawbacks. Obtaining fibroblasts from the skin is an invasive and painful procedure. As well, dermal fibroblasts exhibit slow turnover and renewal rates and may potentially have accumulated environmentally-associated mitochondrial DNA mutations (such as those caused by ultraviolet radiation exposure from the sun) that may not reflect the underlying biology of the patient^14^. In contrast, obtaining peripheral blood mononuclear cells (PBMCs) from whole blood is a much easier and less invasive means of obtaining biological samples from patients. Due to their rapid turnover and self-renewal rates, constant circulation throughout the body, and lifetime immunological memory^15^, COs generated from PBMCs may offer the advantage of more accurately representing the current disease state of the patient.

Here, we developed a human-derived CO model that allows for the assessment of mitochondrial health at the primary, reprogrammed, and differentiated stages, using iPSCs derived from PBMCs and compared and validated these by comparison to COs derived from human embryonic stem cells (hESCs; Fig. 1, Schema). Characterization of fully functional and preserved mitochondrial health throughout the differentiation to COs is crucial to moving toward understanding their role in brain development and disease. The use of PBMCs is flexible and straightforward compared to the use of fibroblasts obtained from invasive biopsies. We expect our approach to be a starting point for more sophisticated patient-derived brain models to investigate mitochondrial health and neuronal activity in a wide range of human diseases – a way forward in developing a standard of care for mitochondrial medicine.

**Fig. 1.**
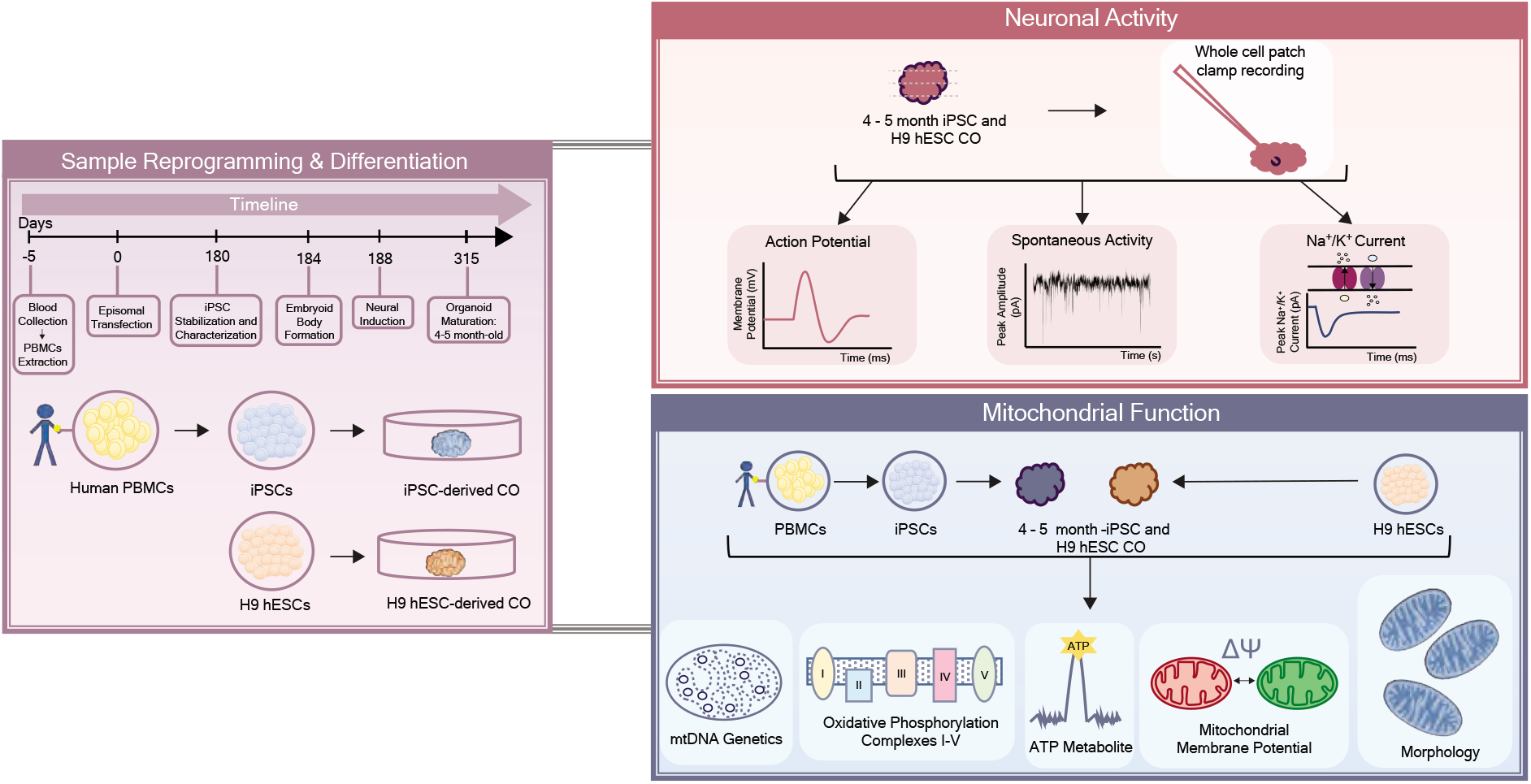
Schematic summary of the study design. Purple panel: An overview and a timeline of sample reprogramming and differentiation from peripheral blood mononuclear cells (PBMCs) to induced pluripotent stem cells (iPSCs) to cerebral organoids (COs) or H9 human embryonic stem cells (H9 hESCs) to COs. Red panel: An overview of electrophysiology experiments (action potentials, spontaneous activity, and sodium and potassium currents) in cerebral organoids. Blue panel: An overview of mitochondrial (mt-) genetics (mtDNA haplogroup, heteroplasmy and copy number), function (oxidative phosphorylation, ATP production and mitochondrial membrane potential) and morphology assessment across PBMCs to iPSCs to COs or H9 hESCs to COs.

## RESULTS

### Generation of cerebral organoids using iPSCs derived from PBMCs

To determine whether a blood sample can be used to make COs (Fig. 2A), we first collected whole blood from a healthy female subject (Table S1 for clinical characteristics). PBMCs from whole blood were isolated using Ficoll density centrifugation and electroporated with episomal vectors expressing five reprogramming factors (Oct4, Sox2, Klf4, L-Myc and Lin28, Fig. 2A and 2B-i). An episomal method was chosen because it has been proven to be the most efficient method for generating integration-free human iPSCs from the blood^16^. We confirmed an apparently normal human, female karyotype in the resulting iPSCs (Fig. 2B-ii), indicating that genetic characteristics were retained during the reprogramming process and corresponded to the parental PBMC lineage. Immunofluorescence staining of iPSCs showed positive expression of key pluripotent proteins (Figs. 2B-iii and iv). At the mRNA level, pluripotent markers in the iPSCs were similar to those observed in H9 hESCs (Fig. 2B-v and Table S2), further demonstrating successful generation of iPSCs from PBMCs.

**Fig. 2.**
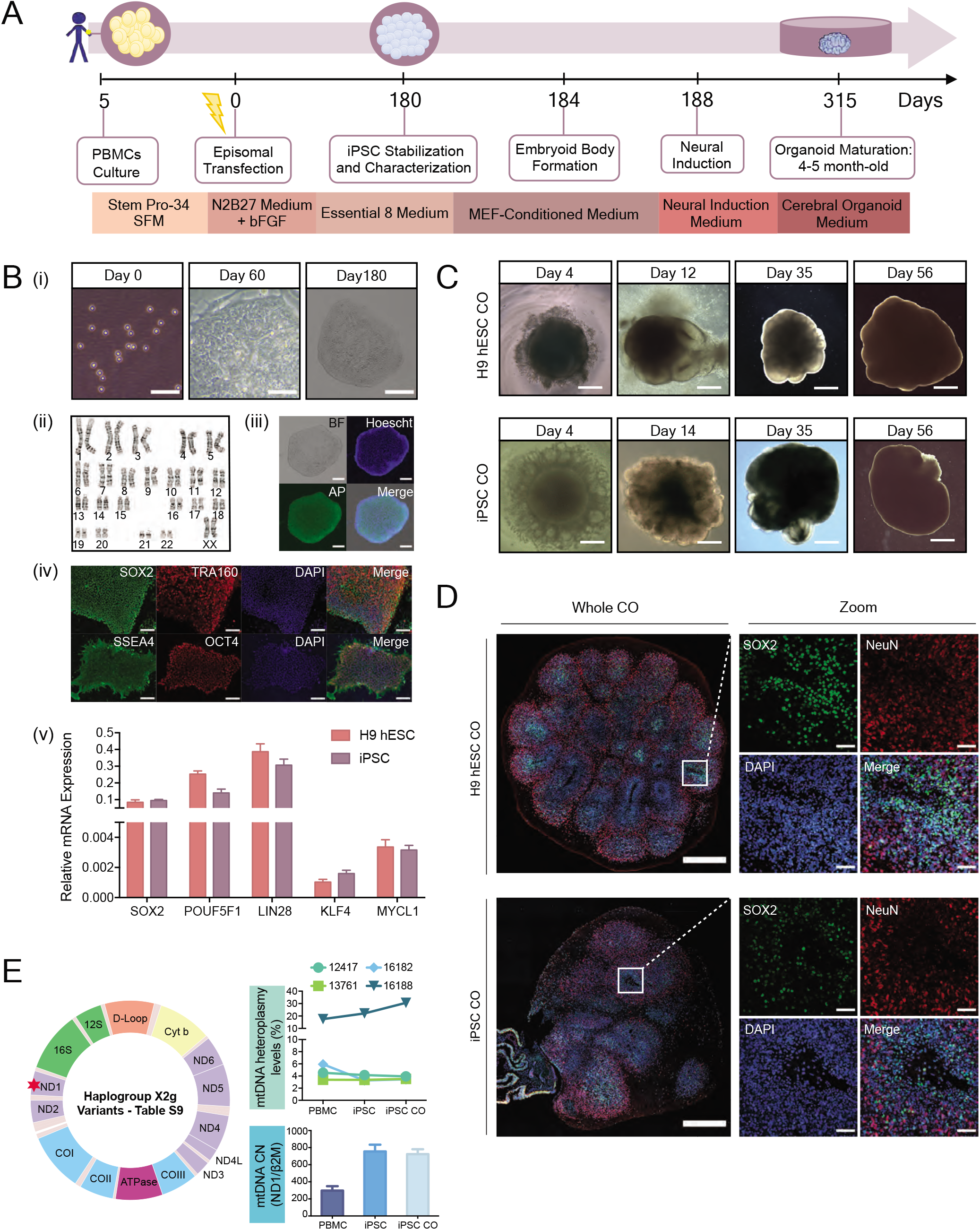
Generation and characterization of cerebral organoids from PBMCs-derived iPSCs and H9 hESCs. **(A)** Timeline and protocol schematic from peripheral blood mononuclear cells (PBMCs) to induced pluripotent stem cells (iPSCs) to cerebral organoids (COs). **(B)** (i) Representative bright-field images showing PBMCs on Day 0, scale bar, 50 μm, reprogramming iPSCs on Day 60, scale bar, 50 μm, and a fully stabilized iPSC colony on Day 180, scale bar, 200 μm; (ii) chromosome analysis with normal female karyotype (46, XX) in 20 cells examined; (iii) representative fluorescent images of iPSCs stained positive for alkaline phosphatase, scale bar, 100 μm; (iv) representative immunofluorescent images of iPSCs stained positive for a set of pluripotency markers, SOX2, TRA160, SSEA4 and OCT4, scale bar, 600 μm; (v) Bar graph showing relative mRNA expression of pluripotency markers, *SOX2, POUF51, LIN28, KLF4* and *MYCL1.* Bars, mean ± SD. **(C)** Representative bright field images showing the progression of cerebral organoid development in H9 hESCs (top panel, scale bars, 250 μm, 250 μm, 500 μm and 1 mm from left to right) and iPSCs (bottom panel, scale bars, 250 μm, 500 μm, 500 μm and 1 mm from left to right). **(D)** Representative fluorescent immunohistochemistry images of whole cerebral organoid in 4.5-month H9 hESC CO and iPSC CO section view (scale bar, 500 μm) and magnified view (scale bar, 50 μm) expressing nuclei (DAPI), radial glia (SOX2) and mature postmitotic neurons (NeuN). Note that immunofluorescent images were used for qualitative observations. **(E)** Mitochondrial (mt-) DNA haplogroup, heteroplasmy and copy number (CN) characterization across PBMCs, iPSCs and iPSC COs. Top right: graph showing mtDNA heteroplasmy levels at four nucleotide positions (MT-12417, MT-13761, MT16182 and MT-16188). Bottom right: graph showing relative mtDNA CN, expressed as MT-ND1/β2M ratio. Schematic of the mitochondrial DNA (left). Red star denotes the mtDNA region used to evaluate mtDNA CN. For more information on the haplogroup X2g variants, see Table S9.

We previously established a robust protocol that allows for the reproducible production of COs from human pluripotent stem cells^17^. These COs display consistent cell type composition and proportions across different batches, making this CO platform useful for disease research as it addresses the problem of variability^17^. Using this protocol, we generated 4.5-month old COs from PBMCs-derived iPSCs (N=14, Table S3) and demonstrated that they have similar overall morphology as those derived from the H9 hESCs (N=14, Fig. 2C and Fig. S1-A). Immunofluorescence staining of the histological sections showed the presence of radial glia (SOX2) and mature neuronal (NeuN) cell types in iPSC-derived COs, which were also present in H9 hESC-derived COs (Fig. 2D and Table S4 for replicates). These qualitative imaging results confirm successful *in vitro* differentiation of iPSCs derived from human PBMCs into COs. As a routine quality control to monitor the genomic integrity across the production stages (PBMCs, iPSC, iPSC-derived CO), we performed a simple sex characterization experiment and demonstrated no change to the female genomic DNA (Fig. S1-B, Table S5 and Table S6), allowing us to use these COs for downstream investigation.

### Electrophysiological responses in iPSC-derived cerebral organoids

A major functional test of any neuronal preparation is the ability to form functional mature neurons with active synaptic neurotransmission^18^. These properties can be tested using electrophysiological recordings from individual neurons and analysis of their action potential (AP)-generation and synaptic activity. Whole-cell recordings were performed in acute slices prepared from COs derived from iPSCs and H9 hESCs (Fig. 3 and Table S4 for replicates). All recorded neurons (N=26 cells for H9 hESC-derived COs and N=29 cells for iPSC-derived COs) were divided into three types based on a combination of electrophysiological properties. Type 1 neurons were immature, could not generate APs, had higher membrane resistance and lower capacitance (Table S7), and relatively small sodium and potassium currents triggered by depolarization (Fig. 3B). Spontaneous activity was not recorded in type 1 neurons. Both type 2 and 3 neurons were able to generate APs; however, type 2 neurons had smaller amplitude and slower kinetics when compared to type 3 neurons (Fig. 3A). Correspondingly, sodium and potassium currents were smaller in type 2 neurons compared to type 3 (Fig. 3B). In addition, only mature, type 3 neurons generated stable trains of spontaneous APs at holding membrane level or during slight depolarization (Fig. 3C). However, spontaneous AP frequency and synaptic activity were similar between type 2 and 3 neurons (Fig. 3C).

**Fig. 3.**
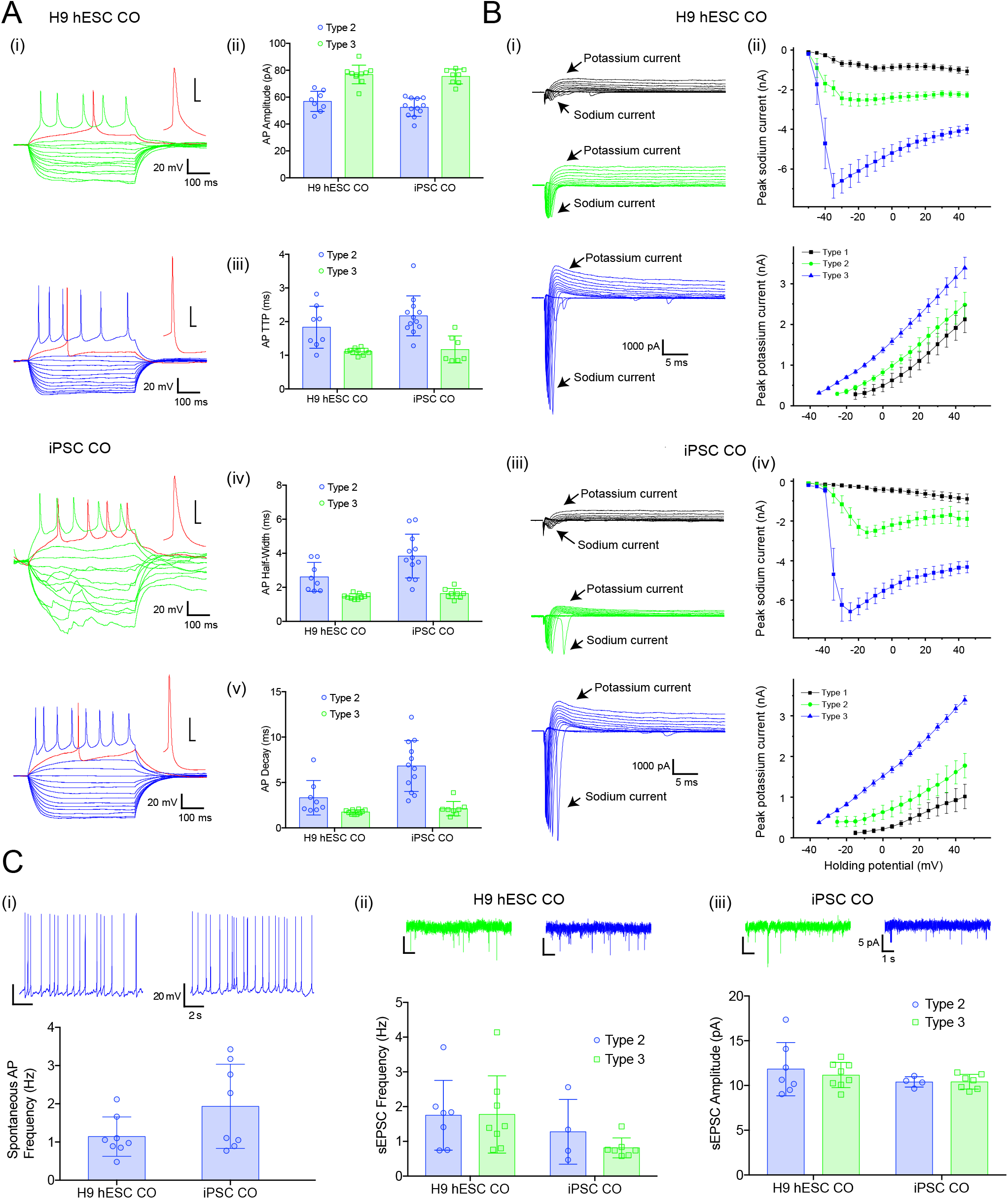
Electrophysiological characterization in H9 hESC-derived COs and iPSC-derived COs. **(A)** (i) Examples of voltage responses to hyperpolarizing and depolarizing set of currents recorded from type 2 (green) and type 3 (blue) neurons. Individual APs are expanded in the insert (red), scale 20 mV/ 5 ms. Summary bar graphs showing (ii) AP amplitudes, (iii) rise time, (iv) half-width and (v) decay, note that APs have lower amplitude and slower kinetic in type 2 neurons. **(B)** (i) Examples of inward sodium and outward potassium currents recorded from neurons type 1 (black), type 2 (green) and type 3 (blue). Scale bar is the same for all traces. Summary current-voltage plots showing increase in peak current amplitude for both sodium (top) and potassium (bottom) during neuronal maturation (ii and iii). These plots look similar independently of COs origin. **(C)** (i) Examples of spontaneous AP firing in type 3 neurons (top) and summary bar graph of AP frequency in H9 hESC- and iPSC-derived COs (bottom). (ii) Examples of spontaneous excitatory postsynaptic currents (EPSCs) recorded from type 2 (green) and type 3 (blue) neurons. Scale bar is the same for all traces. (iii) Summary bar graphs showing that spontaneous synaptic activity including sEPSC frequency (left) and amplitude (right) was similar in both types of neurons regardless origin of COs. Error bars show standard deviation.

Next, the electrophysiological properties of type 2 and 3 neurons in COs generated from PBMC-derived iPSCs were compared to those found in H9 hESC-derived COs. We found similar AP amplitude (Fig 3A-ii) and rise time (Fig. 3A-iii) in type 2 neurons between the two types of COs. Type 2 neurons from iPSC-derived COs showed slower half-width (Fig. 3A-iv) and decay (Fig. 3A-v) while type 3 mature neurons showed no difference in any measured AP properties (Figs. 3A-i to v). In addition, the amplitude of sodium or potassium currents in type 2 and 3 neurons were similar between both types of COs (Fig. 3B). Finally, we showed similarities in spontaneous synaptic activity of type 2 or type 3 neurons (Figs. 3C-ii and iii), and spontaneous AP frequency in mature type 3 neurons (Fig. 3C-i), indicating that iPSC-derived COs contain mature neurons with electrophysiological properties similar to H9 hESC-derived COs.

### Mitochondrial genetics, function and morphology from PBMCs, iPSCs to COs

Mitochondrial function has been classified as a key indicator of neuronal activity and healthy cells. Thus, we next monitored the integrity and stability of mitochondrial genotype, function and morphology across the primary (PBMCs), reprogrammed (iPSCs), and differentiated (COs) stages (Table S4 for replicates).

Mitochondria harbours its own genome in multiple copies, called the mitochondrial DNA (mtDNA). The mtDNA encodes required subunits of the electron transport and oxidative phosphorylation complexes as well as the ribosomal and transfer RNAs required for their translation^19^. As a result, it is critical to evaluate whether mtDNA integrity is preserved throughout the process of generating COs^20^. The integrity of mtDNA can be evaluated by identifying the: 1) haplogroup, which is defined by a set of genetic variants associated with maternal ancestry; 2) percentage of heteroplasmy (mix of normal and mutated mtDNA) or homoplasmy (uniform collection of mtDNA, either mutated or normal) and; 3) copy number.

Using mtDNA sequencing, we identified the X2g haplogroup across PBMCs, iPSCs and iPSC-derived COs (Fig. 2E, Table S8, Table S9). We further validated these results by performing polymerase chain reaction (PCR) amplification of short mtDNA fragments corresponding to the X2g haplogroup and sequencing these products (Table S10). No change to the mtDNA haplogroup was observed (Table S9), confirming that the mitochondrial ancestry of the donor was retained throughout the process of CO production. The mtDNA also lacks repair mechanisms, making it susceptible to changes caused by the culture environments that could potentially lead to novel mutations in the iPSCs and COs. To ensure the preservation of mtDNA integrity throughout the CO production, we evaluated the complete mtDNA sequence and confirmed that the nucleotide identity across PBMCs, iPSCs and iPSC-derived COs is 100%. As no novel mutations were introduced, these results assure that we have retained the mtDNA identity of the donor during the reprogramming and differentiation process. A recurring question in the scientific community is whether there is a selection process that drives homoplasmy from heteroplasmy upon iPSC reprogramming^21–23^. Using the mtDNA sequencing data, we demonstrated the conservation of four heteroplasmic variants across PBMCs, iPSCs and iPSC-derived COs suggesting no selection process occurred (Fig. 2E). To further evaluate mtDNA integrity, we performed quantitative polymerase chain reaction (qPCR) using specific primers that target a highly conserved gene in the mtDNA (NADH dehydrogenase subunit 1, MT-ND1) and quantified the ratio of MT-ND1 gene copy to a single-copy nuclear gene (Beta-2-Microglobulin, b2M). Our results revealed multiple copies of mtDNA, which were preserved upon CO production from iPSCs (Fig. 2E, Table S11). Altogether, we provided solid evidence that the iPSC-derived COs retained the mtDNA genetic identity of their somatic origin (PBMC), resulting in the recapitulation of the donor’s mitochondrial phenotype.

While mtDNA integrity was preserved, a remaining question is whether this translates to healthy and functional mitochondria. As a first approach, we stained COs before and after differentiation using MitoTracker Red CMXRos, which is a cationic fluorescent dye that penetrates solely into active and live mitochondria in a potential-dependent manner. Following sample staining, we demonstrated MitoTracker-positive cells across PBMCs, iPSCs and iPSC-derived COs, which were similarly detected in H9 hESCs and H9 hESC-derived COs (Fig. 4A). As the mitochondria appeared live and active, we next investigated function and responsiveness. The functional state of the mitochondria can be evaluated by monitoring changes in the mitochondrial membrane potential (MMP)^24^. JC-1 (5,5′,6,6′-tetrachloro-1,1′,3,3′-tetraethylbenzimi-dazolylcarbocyanine iodide) is a dual emission cationic dye that accumulates in the mitochondria in response to the MMP, yielding a green fluorescence in less energetic mitochondria and red fluorescent aggregates in highly energetic mitochondria. Here, we used JC-1 dye as it is optimal for the end-point analysis of mitochondrial health^25^. Treatment of the H9 hESC- and iPSC-derived COs before and after differentiation with JC-1 led to an increased red-to-green fluorescence ratio, indicative of higher MMP, corresponding to a highly energetic mitochondrial state (Fig. 4B and Fig. S2-A). We next investigated whether these COs, both before and after differentiation, remained responsive to pharmacological intervention. For this, we administered carbonyl cyanide-*4*-(trifluoromethoxy)phenylhydrazone (FCCP), an uncoupler of oxidative phosphorylation (OXPHOS) that collapses and depolarizes the MMP. Treatment with FCCP led to a marked decrease in the red-to-green fluorescence ratio and fragmentation of the mitochondrial tubular network into smaller circular monomers (Fig. 4C and Fig. S2-A), indicating that in all cases, the COs are responsive to pharmacological intervention. Overall, these data demonstrate that the mitochondria remained functional and responsive throughout the process of generating COs.

**Fig. 4.**
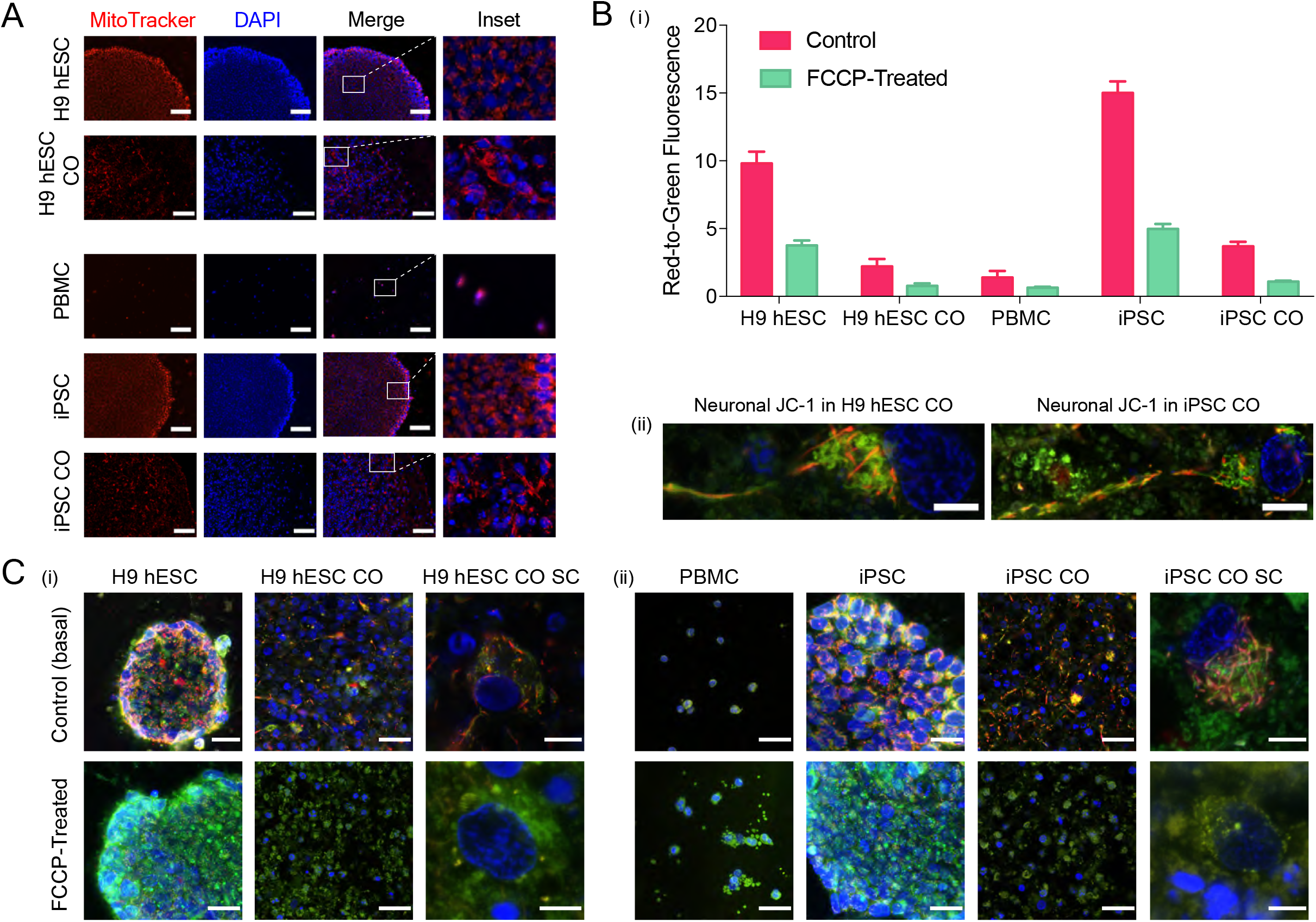
Characterization of active, functional and responsive mitochondria throughout CO generation. **(A)** Active mitochondria in H9 hESCs (scale bar, 100 μm), H9 hESC COs (scale bar, 50 μm), PBMCs (scale bar, 100 μm), iPSCs (scale bar, 100 μm) and iPSC COs (scale bar, 65 μm) stained with MitoTracker Red CMXRos (red) and DAPI (nuclei, blue). Last column shows insets, enlarged views of boxed areas from the merge images. **(B)** (i) Bar graph summarizing mitochondrial membrane potential (MMP) as red-to-green fluorescence ratio across primary, reprogrammed and differentiated stages in samples treated with JC-1 only (control basal MMP level) and those treated with carbonyl cyanide-4-(trifluoromethoxy)phenylhydrazone (FCCP). Fluorescence was recorded using a microplate reader. Bars, mean ± SD; (ii) Representative MMP images of a fully captured neuron with cell body and axon projection in H9 hESC CO (left, scale bar, 5 μm) and iPSC CO (right, scale bar, 7 μm). Red, JC-1 aggregates (highly energetic or polarized), green, JC-1 monomers (less energetic or depolarized). **(C)** (i) Top row shows representative MMP images across CO generation from H9 hESCs (scale bar, 25 μm) to H9 hESC CO (scale bar, 25 μm) or hESC CO single cell (SC, scale bar, 10 μm) and; (ii) from PBMCs (scale bar, 25 μm) to iPSCs (scale bar, 25 μm) to iPSC COs (scale bar, 25 μm) or iPSC CO SC (scale bar, 5 μm). Bottom row shows representative images of samples treated with FCCP followed by JC-1 to visualize the collapse of MMP in (i) H9 hESCs (scale bar, 25 μm) to H9 hESC CO (scale bar, 25 μm) or hESC CO SC (scale bar, 5 μm) and; (ii) from PBMCs (scale bar, 25 μm) to iPSCs (scale bar, 25 μm) to iPSC COs (scale bar, 25 μm) or iPSC CO SC (scale bar, 5 μm). All images were captured at 100x magnification with slight variations in the zoom factor to capture the single cell(s) or single colony of interest. Immunofluorescent images shown here were used for qualitative observations only.

The MMP is generated upon the passing of electrons through a group of mitochondrial protein complexes I, III, and IV located on the inner mitochondrial membrane, which in turn is used to drive adenosine 5′-triphosphate (ATP) production through complex V. We determined the ability of COs derived from iPSCs and H9 hESCs to perform OXPHOS by measuring the assembly levels of intact mitochondrial complexes and the intracellular ATP levels. Using Luminex bead-based multiplex immunoassay, we successfully tracked the expression of intact mitochondrial complexes I-V across PBMC, iPSC and iPSC-derived COs and H9 hESC and H9 hESC-derived COs (Fig. 5A). To visualize the proper formation and distribution of OXPHOS, we performed immunofluorescence staining of TOMM-20, a marker of the outer mitochondrial membrane and NDUFS3, SDHA, UQCRC1, COX-IV, and ATP Synthase-β, markers of the inner mitochondrial membrane of complexes I, II, III, IV and V, respectively, and observed expression and typical distribution of mitochondria throughout the COs and cells (Fig. 5B and Table S2). To evaluate intracellular ATP production, we performed CellTiter-Glo luminescent assay and shown the maintenance of ATP production throughout the generation of COs, which was in a similar pattern to that observed for the OXPHOS data indicating a high production of ATP in stem cells as compared to COs (Fig. 5C and Fig. S2-B). Energy metabolism has been suggested to play a crucial role in the regulation and maintenance of pluripotency^26-28^. To examine this further, we used oligomycin, an inhibitor of complex V, to block ATP production by mitochondrial OXPHOS in PBMC-derived iPSCs and H9 hESCs. We observed a subtle decrease (~20%) in ATP production (Fig. S2-C), suggesting that the immediate source of ATP production is glycolysis in iPSCs and H9 hESCs. Although less efficient in terms of energy production, stem cells have been shown to use this pathway at a much faster rate which is crucial for maintaining pluripotency and may explain the high ATP production^26,27,29^.

**Fig. 5.**
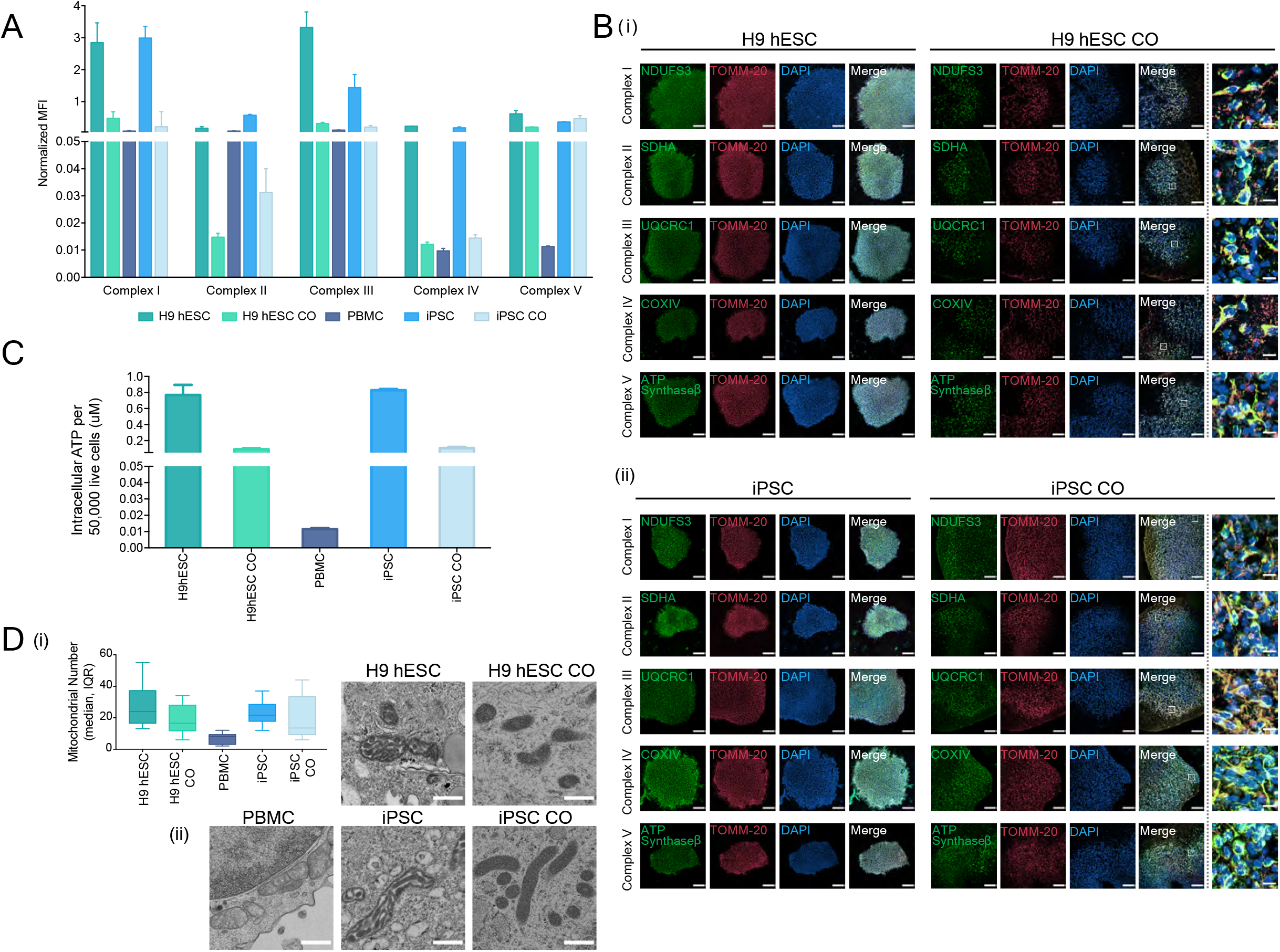
Mitochondrial oxidative phosphorylation and morphology throughout CO generation. **(A)** Bar graph showing the oxidative phosphorylation (OXPHOS) complexes I – V assembly levels in H9 hESCs, H9 hESC COs, PBMCs, iPSCs and iPSC COs. Median fluorescence intensities for each complex were recorded using Luminex technology. Bars, median fluorescence intensity ± SD. **(B)** Representative immunofluorescence images showing the formation of the inner mitochondrial membrane OXPHOS proteins (complex I, NDUFS3; complex II, SDHA; complex III, UQCRC1; complex IV, COXIV; complex V, ATP synthase-β; green) and the outer mitochondrial membrane (TOMM-20; red) in (i) H9 hESCs to H9 hESC COs and; (ii) iPSCs to iPSC COs. All scale bars, 100 μm. Last column shows insets, enlarged views of boxed areas from the merge CO images, all scale bars, 10 μm. In the CO tissues, out of focus light and autofluorescence of tissue matrix led to background noise which were corrected for better visualization. Brightness levels of each images were also adjusted to optimize visualization. Immunofluorescent images shown here were used for qualitative observations only - no quantitative analyses were performed. **C)** Bar graph showing the intracellular ATP levels across H9 hESCs to H9 hESC COs and PBMCs, iPSCs to iPSC COs, bars, mean ± SD. **(D)** (i) Box plot summarizing the median number of mitochondria in 10 cells examined, boxes, median and interquartile range, IQR. (ii) Representative electron micrographs of mitochondrial morphology across H9 hESCs (scale bar, 5 μm) to H9 hESC COs (scale bar, 500 nm) and PBMCs (scale bar, 5 μm), iPSCs (scale bar, 500 nm) to iPSC COs (scale bar, 500 nm).

Lastly, we monitored mitochondrial number and visualized morphology using transmission electron microscopy (TEM). Consistent with OXPHOS and ATP production, H9 hESCs and iPSCs had higher number of mitochondria compared to their corresponding COs (Fig. 5D-i). Moreover, the electron micrographs revealed well-preserved mitochondrial morphologies with defined cristae before and after the generation of COs (Fig. 5D-ii). Consistent with expectations, mitochondria in the iPSCs and H9 hESCs displayed globular and immature shapes with electron-dense cristae whereas more elongated and mature mitochondria with thinner cristae were visually observed in the COs (Fig. 5D-ii), suggesting mitochondrial maturation during the differentiation process.

## DISCUSSION

To the best of our knowledge, this is the first report demonstrating the development of a functional human-derived CO model made from PBMCs successfully tracking overall mitochondrial health, including genetics, function and morphology. Although the COs are immature, more closely resembling a fetal rather than an adult brain, they exhibited electrophysiological responses including both mature and immature neurons and displayed functional and responsive mitochondria across all stages of development, from PBMCs to iPSCs to iPSC-derived COs. We demonstrated the preservation of the donor’s mtDNA genetic integrity, highlighting the potential of COs generated from PBMC-derived iPSCs to recapitulate the functional phenotypes that may be rooted in their genetic information. We further demonstrated the response of iPSC-derived COs to pharmacological treatment (FCCP), mimicking the individual’s overall cellular/mitochondrial response, which supports the use of iPSC-derived COs as a representative model to evaluate function and to screen for therapeutic compounds.

Despite encouraging progress, CO models do have limitations. An ongoing challenge is that with prolonged culture, COs reach a size limit along with the occurrence of cell death in the centre regions as a result of insufficient diffusion of oxygen and nutrients^11,30^. While it may be possible to increase oxygen availability by growing COs in a higher oxygen environment, this can be toxic if not carefully controlled^31^. So far, COs do not develop vasculature that could provide nutrients, but studies involving vascularization and generation of blood vessel organoids are underway and have yielded some promising results^32,33^. The fetal nature of these COs is also a limitation to be noted as current methods do not allow for the full development of COs into the adult brain. Taking these limitations into consideration, any interpretation of results requires caution when considering CO models.

While acknowledging that CO models currently do not have the full precision of a human brain, the ability to interrogate overall mitochondrial health in a human context is extremely valuable for studying brain development and diseases. By evaluating mitochondrial health in both the PBMCs and iPSC-derived COs, we can start to identify changes in the brain that may also be present in the blood of patients. As a result, this approach can aid in the development of biomarkers, companion diagnostics, and novel mitochondrial therapeutics that can inform critical clinical decisions in mitochondrial medicine. As there is currently no standard of care for the assessment of mitochondrial health in both the brain and periphery of patients, the platform provided here is a potential starting point that can be applied to a wide range of disease phenotypes.

## MATERIALS AND METHODS

### Blood sample collection and processing

We followed the guidelines established by the Biomarkers Task Force as modified by the World Federation of Societies of Biological Psychiatry for clinical assessment and documentation, ethical procedures and blood sample collection^34^. This study was performed in accordance with the latest version of the Declaration of Helsinki and approved by the Research Ethics Board at the University of Toronto, Canada (Protocol Number: 29949). Venous blood (10 mL) was drawn from a participant through venipuncture into a plastic whole-blood tube with spray-coated K2EDTA (16 x 100 mm x 10.0 mL BD Vacutainer^®^ Plus). Whole blood was carefully layered on Ficoll-Paque (GE Healthcare, 17144002) at a 1:1 ratio in a conical tube and centrifuged at 400 x g for 40 minutes. Following centrifugation, peripheral blood mononuclear cells (PBMCs) from the buffy coat were isolated, washed twice with Dulbecco’s phosphate-buffered saline (DPBS) (Gibco™, 14190250), and immediately seeded in a 25 cm^2^ tissue culture flask at a density of about 2 x 10^6^ cells containing 10 mL of StemPro^®^-34 SFM (Gibco™, 10639011) supplemented with GlutaMAX-I (1X), recombinant human SCF (100 ng/mL), recombinant human IL-3 (50 ng/mL), and recombinant human GM-CSF (25 ng/mL) for 72 hours at 37°C with 5% CO_2_.

### Generation of induced pluripotent stem cells

PBMCs were transfected with Epi5™ EBNA-1/OriP-based episomal vectors containing five pluripotent factors (Oct4, Sox2, Klf4, L-myc and Lin28) using the Neon^®^ Transfection System (Invitrogen™, A15960) following the manufacturer’s protocol. Briefly, PBMCs at a density of 2 x 10^5^ per mL were suspended in the buffer supplied by the manufacturer. Equal amounts (1 μg) of Epi5™ reprogramming and the Epi5™ p53-EBNA vectors were used to transfect the PBMCs via electroporation by setting the parameters to 1650 V, 10 ms and 3 pulses. Following transfection, PBMCs were distributed into preconditioned plates coated with 1:50 Geltrex™ LDEV-Free, hESC-Qualified, Reduced Growth Factor Basement Membrane Matrix (Gibco™, A1413302) containing StemPro^®^-34 SFM (Gibco™, 10639011) media supplemented with GlutaMAX-I (1X), recombinant human SCF (100 ng/mL), recombinant human IL-3 (50 ng/mL), and recombinant human GM-CSF (25 ng/mL). After 24 hours of incubation at 37°C with 5% CO_2_, cells were transitioned to N2B27 media, composed of DMEM/F-12, HEPES, N-2 Supplement (1X), B-27™ Supplement (1X), MEM Non-Essential Amino Acids Solution (1X), GlutaMAX-I (0.5X), 2-Mercaptoethanol (1:500), FGF-Basic (AA 1-155) Recombinant Human Protein (100 ng/mL). N2B27 media was changed every other day with fresh FGF-Basic (100 ng/mL) until emergence of reprogramming induced pluripotent stem cells (iPSC) colonies. iPSC colonies that exhibited embryonic stem cell (ESC)-like morphologies were picked under a stereomicroscope for expansion and maintenance in Essential 8™ medium (Gibco™, A1517001).

### Characterization of induced pluripotent stem cells

We followed the protocol developed by Marti et al. (2013) showing the different steps required to characterize an iPSC line, which includes: 1) pluripotency test (alkaline phosphatase and immunodetection of pluripotency markers) and 2) karyotype^35^.

#### Alkaline phosphatase live stain

iPSCs were stained with Alkaline Phosphatase (AP) Live Stain (Invitrogen™, A14353) as per the manufacturer’s protocol. iPSCs cultures were incubated with AP Live Stain diluted at 1:500 in DMEM/F-12, HEPES media for 30 minutes at 37°C with 5% CO_2_ and 2 drops/mL of NucBlue™ Live ReadyProbes™ Reagent (Invitrogen™, R37605) for 20 minutes. Following staining, cells were washed twice with DMEM/F-12, HEPES media (5 minutes per wash) to remove excess substrate. After the final wash, FluoroBrite™ DMEM (Gibco™, A1896701) media was added and iPSCs were immediately visualized using the EVOS FL Cell Imaging System (Invitrogen™).

#### Detection of pluripotency markers

Protein expression of pluripotent markers (SSEA4, OCT4, SOX2, TRA-1-60) in iPSCs was performed using the Pluripotent Stem Cell 4-Marker Immunocytochemistry Kit (Invitrogen™, A24881). iPSC colonies were fixed for 15 minutes, permeablized for 15 minutes, and blocked for 30 minutes at room temperature. Two primary antibody combinations were used to incubate the iPSCs: antibody combination 1: anti-SSEA4 and anti-OCT4; and antibody combination 2: anti-SOX2 and anti-TRA-1-60 for 3 h at room temperature. After washing, iPSCs were incubated with secondary antibodies (Alexa Fluor^®^ 594 donkey anti-rabbit for anti-OCT4, Alexa Fluor^®^ 488 goat anti-mouse IgG3 for anti-SSEA4, Alexa Fluor^®^ 488 donkey anti-rat for anti-SOX2 and Alexa Fluor^®^ 594 goat anti-mouse IgM for anti-TRA-1-60) for 1 h. Following wash, 1 drop of NucBlue fixed cell stain (DAPI; Invitrogen) was added, and images of iPSCs were captured using a Nikon Eclipse Epifluorescent microscope equipped with a 12-bit Q-imaging camera, Retiga 1300B (7323). To validate pluripotency at the messenger RNA (mRNA) level, we performed quantitative reverse transcription polymerase chain reaction (qRT-PCR). The experiment was performed using wet bench verified predesigned RT^2^ qPCR primers on SOX2, POUF5F1, KLF4, MYCL1, LIN28A, GAPDH (control). Total RNA was extracted from the iPSCs and H9-hESCs using the RNeasy Mini Kit (Qiagen) as per the manufacturer’s protocol. The cDNA was synthesized from 500 ng of total RNA following the Qiagen’s RT^2^ First Strand Kit. Each PCR reaction contained 1 μL cDNA mixture, 1 μL RT^2^ qPCR primers (SOX2 or POUF5F1 or KLF4 or MYCL1 or LIN28A or GAPDH), 10.5 μL of water, and 12.5 μL of RT^2^ SYBR Green master mix. After 46 cycles of amplification, data was acquired using BioRad CFX96 RT-PCR detection system with the following cycling conditions: 1 cycle of 10 min at 95°C to activate HotStart DNA Taq Polymerase and 45 cycles of 15 s at 95°C, followed by 1 min at 60°C for fluorescence data collection. The expression level of each gene of interest was normalized to the housekeeping gene and was expressed as mean 2^-[CT (gene of interest) – CT(GAPDH)]^.

#### Karyotype

Karyotype analysis via G-banding was performed on cells from two 25 cm^2^ tissue culture flasks submitted to the TCAG Cytogenomics Facility. When cells reached 50% confluence, Karyomax Colcemid^®^ was added to each flask to a final concentration of 0.15 μg/mL (Cat. #15212-012, Gibco, CA, USA) and incubated in 37°C CO_2_ incubator for 1 hour for one flask, and 30 minutes for the second flask. Cells were then collected by treating with 1 mL 0.05% trypsin, 0.53mM EDTA (Cat. #325042 EL, WISENT Inc., Quebec, Canada) at 37°C for 5 minutes, and pipetted up and down five times to break into single cells. Cells from each flask were centrifuged in a 15mL conical tube at 1000 rpm for 10 minutes, supernatant was aspirated until 0.5 mL remained, and cell pellet was flicked to resuspend completely. Cells were suspended in 8 mL buffered hypotonic solution (0.054M KCl, 0.02% EGTA, 20mM HEPES, pH 7.4), and incubated in 37°C for 25 minutes. Eight drops of Carnoy’s fixative (methanol/acetic acid, 3:1) were added and mixed together. Cells were centrifuged at 1000 rpm for 10 minutes, supernatant was aspirated until 0.5 mL remained, and cell pellet was flicked to resuspend completely. Carnoy’s fixative was added to the 14 mL mark, tubes were inverted to mix, and incubated in −20°C for at least one hour to overnight. After two more rounds of fixations (add 8 mL fixative, invert to mix, and centrifuge at 1000 rpm for 10 minutes), cells were resuspended in 0.5-1 mL of fixative and cells from each suspension were dispensed onto glass slides and baked at 90°C for 1.5 hours, followed by overnight aging at room temperature in desiccator. Routine G-banding analysis was then carried out. Twenty metaphases per cell line were examined.

### Generation of cerebral organoids from H9 hESCs and iPSCs

Cerebral organoids were generated from iPSC and H9 hESCs as previously described^17^. In brief, stem cells were aggregated into embryoid bodies and after induction of the neural germ layer were embedded in Matrigel droplets in differentiation media. Once removed from the Matrigel, free-floating organoids were maintained in prolonged culture on an orbital shaker.

### Fluorescent immunohistochemistry for the characterization of cerebral organoids

Whole COs derived from H9 hESCs and iPSCs were fixed with 4% paraformaldehyde at 4°C overnight on an orbital shaker, cryoprotected in 30% (weight/volume) sucrose at 4°C overnight and embedded in optimal cutting temperature compound. Cryosections of 14 μm thickness were collected on Superfrost Plus Microscope glass slides (Fisher Cat# 12-550-15) using a Leica cryostat and were stored in −80°C until analysis. Prior to immunofluorescent staining, cryoslides from H9 hESCs- and iPSCs-derived COs were equilibrated to room temperature and tissues were rehydrated with 0.1% Tween-20 in 1X PBS pH 7.4. Tissues were then incubated in permeabilization/blocking solution (1X PBS pH 7.4, 10% Normal Donkey Serum, 2% BSA, 0.5% Triton X-100) for 1 hour at RT in a humidified chamber. Proteins of interest were labeled with the following primary antibodies: SOX2 (goat, R&D, AF2018, 3 μg/mL) and NeuN (rabbit, Cell Signaling Technology, 12943S, 1:500) diluted in antibody blocking solution (1X PBS pH 7.4, 0.1% Tween-20, 1% BSA, and 5% Normal Donkey Serum) at 4°C and incubated overnight in a humidified chamber. Following overnight incubation, tissues were washed three times (5 minutes per wash) with 0.1% Tween-20 in 1X PBS pH 7.4 and then incubated overnight at 4°C with the appropriate secondary antibodies: Donkey anti-Goat IgG (H+L) Highly Cross-Adsorbed Secondary Antibody Alexa Fluor Plus 555 (Invitrogen, A32816) and Donkey anti-Rabbit IgG (H+L) Highly Cross-Adsorbed Secondary Antibody Alexa Fluor Plus 647 (Invitrogen, A32795) diluted 1:750 in antibody blocking solution. Following secondary antibody incubation, tissues were washed three times in 1X PBS pH 7.4 with 0.1% Tween-20. Nuclei were counterstained by adding a small drop of ProLong™ Gold Antifade Mountant with DAPI (Invitrogen™, P36935) to each tissue section prior to sealing under glass coverslips. Mountant was left to cure overnight at room temperature and protected from light. Images were captured using Leica TCS SP8 lightning confocal laser scanning microscope (DMI6000) equipped with white light laser (470-670 nm). Z-stack images were taken at 1μm intervals and images were collected using LASX software (Lecia Microsystems). See Table S4 for biological and technical replicates.

### Acute slice preparation

Brain organoids (4.5 months old) were embedded into 4% low melting agarose and 300 μm slices were cut with vibrating-blade microtome (Leica, Germany). Slices were prepared using ice-cold artificial cerebrospinal fluid (ACSF) containing (mM): NaCl 87, NaHCO_3_ 25, KCl 2.5, NaH_2_PO4 1.25, MgCl_2_ 7, CaCl_2_ 0.5, glucose 25 and sucrose 75. Right after sectioning, slices were placed in oxygenated ACSF at room temperature and used for experiments within 4 hours. All recordings were performed with extracellular ACSF containing (mM): NaCl 124, NaHCO_3_ 25, KCl 2.5, MgCl_2_ 1.5, CaCl_2_ 2.5 and glucose 10, equilibrated with 95% O_2_–5% CO_2_, pH7.4, maintained at 29–32°C and perfused at a rate of 2-3 mL/min.

### Electrophysiology

Whole-cell current-clamp and voltage-clamp recordings were made with glass electrodes (3 −5 MΩ) filled with a solution containing (mM): K-gluconate 120, KCl 20, MgCl2 2, EGTA 0.6, MgATP 2, NaGTP 0.3, Hepes 10, phosphocreatine. Individual neurons were visually identified usually around slice edge in order to increase chance of patching mature neurons. Electrophysiological recordings were made using a Multi Clamp 700A amplifier (Axon Instruments, Union City, CA, USA), operating under current-clamp and voltage-clamp mode. Data were filtered at 4 kHz, and acquired using pClamp 10 software (Molecular devices, Sunnyvale, CA, USA). All recordings were done at a holding potential −70 mV. The uncompensated series resistance was monitored by the delivery of −10 mV steps throughout the experiment, only recordings with less than 15% change were analyzed.

Resting membrane potential was measured immediately after establishing whole-cell patch clamp recording and followed by measurements of passive neuronal properties (access resistance, membrane resistance and capacitance) using automatic membrane test in pClamp 10 software Ability to generate spontaneous action potentials was tested in current clamp mode starting from holding membrane potential and by application of depolarizing steps (5 pA increments) until steady firing was reached. This steady firing was observed only in mature neurons. Voltage-dependent sodium and potassium currents were measured in voltage clamp mode, using 500 ms voltage steps applied from a holding potential to a range of potentials between −50 and +50 mV (in 10 mV increments). Action potentials were triggered using 500 ms depolarizing pulses increasing amplitude (in 5pA steps). Synaptic events were analyzed using pClamp 10 software within 5 minutes of recordings, individual events were detected using automatic template search. Templates were created using the average of at least 10 events aligned by the rising of their slopes. AP amplitude, rise time, half width and time to peak (TTP) were calculated to investigate excitability. See Table S4 for biological and technical replicates.

### Total DNA extraction

Genomic DNA was isolated from PBMCs, stem cells (H9 hESCs and iPSCs) and COs derived from H9 hESCs and iPSCs using the QIAamp DNA Mini Kit (Qiagen, 51304) by following the manufacturer’s recommended protocol and using the recommended elution volumes. The concentration, purity and integrity of all extracted DNA was determined by spectrophotometric measurement on the NanoDrop^®^ ND-1000 (Thermo Scientific™).

### Sex and haplogroup characterization

Primer pairs for haplogroup and sex determination (which included a genomic control primer pair) are listed in Figs. 2-5 and 2-8. PCR reactions for sex determination or haplogroup were performed on 5-10 ng genomic DNA in a 25μl reaction mix containing 16.6 mM (NH4)2SO4, 67mM Tris-HCl pH 8.8, 6.7 mM MgCl2, 5 mM 2-Mercaptoethanol, 6.7 mM EDTA, 10% Dimethylsulfoxide, 0.125mg/ml Bovine Serum Albumin (Bioshop), 1 mM each deoxynucleotide triphosphate (Biobasic), 0.5uM each primer (IDT) and 1.5u DreamTaq (Thermofisher). All of the amplifications were performed on a BioRad C1000 thermal cycler. For the sex determination the conditions were 96 °C for 2 min; 12 cycles of 94 °C for 20s, 64-58°C for 30s (reducing 0.5°C per cycle) and 72 °C for 35s, followed by 28 cycles of 94 °C for 20s, 58°C for 30s and 72 °C for 35s with a final extension step at 72 °C for 5 min. For the haplogroup primers, the conditions were 96 °C for 1 min; 40 cycles of 94 °C for 30s, 56°C for 20s and 72 °C for 150s with a final extension step at 72 °C for 5 min. The PCR products for the sex determination were separated on a 1.8% TBE agarose gel containing 0.5 μg/ml Ethidium Bromide at 100 volts and visualized under UV light. Ten PCR products for haplogroup sequencing were purified using Biobasic EZ-10 Spin Column PCR Products Purification Kit following the manufacturer’s instructions and eluted in ddH2O for sequencing with internal primers. Sequencing reactions were performed by The Centre for Applied Genomics (Hospital for Sick Children) on ABI 3730XL machines.

### Mitochondrial DNA sequencing for haplogroup and heteroplasmy characterization

The mitochondrial DNA was amplified as two fragments using two the following two primer sets: 1) COIII-F (5’-TCACAATTCTAATTCTACTGA-3’) and mt16425R (5’-GATATTGATTTCACGGAGGATGGTG-3’) resulting in a 7321bp fragment and 2) mt16426F (5’-CCGCACAAGAGTGCTACTCTCCTC-3’) and COIII-R (5’-CGGATGAAGCAGATAGTGAGG-3’) and 10,011bp products. PCR reactions were performed using TaKaRa LA Taq Hot Start polymerase kit (TaKaRa) and 50ng of total genomic DNA in a 50uL PCR reaction. Cycling conditions included an initial denaturation at 94 °C for 2 min, followed by 30 cycles of denaturation at 94 °C for 20 sec, annealing at 60 °C for 30 sec, and extension at 68 °C for 12 min. The reaction was concluded with a final extension at 68 °C for 20 min. PCR products were analyzed on an agarose gel and HindIII Ladder (New England Biolabs), and subsequently purified with the GeneJet PCR Purification kit (ThermoFisher). Samples were then submitted for Illumina sequencing Nextera XT library prep on a Hiseq 2500 high throughput flowcell 2×125bp.

### Mitochondrial DNA copy number

We adapted a protocol from Picard et al (2015) for measuring mtDNA copy number by qPCR^36^. Briefly, total genomic DNA was extracted as described in the method “Total DNA Extraction” and diluted to 0.1 ng/μL. The ratio of mtDNA to nuclear DNA was quantified by 2^-DCt^ method using the following primer pairs: β-2 microglobulin forward (TGCTGTCTCCATGTTTGATGTATCT) (3′–5′) and reverse (TCTCTGCTCCCCACCTCTAAGT) for nuclear DNA, and ND1 forward (ATGGCCAACCTCCTACTCCT) and reverse (CTACAACGTTGGGGCCTTT) for mtDNA. Each PCR reaction (20 μL) contained 2X SensiFAST SYBR^®^ No-ROX mix (Bioline, BIO-98005, 10 μL), 10μM primers, and 0.6 ng of template DNA. After 40 cycles of amplification, data was acquired using BioRad CFX96 RT-PCR detection system (Bio-Rad Laboratories, Inc.) with the following cycling conditions: 1) 95°C for 3 min; 2) 95°C for 10 seconds and; 3) 60°C for 20 seconds. See Table S4 for biological and technical replicates.

### MitoTracker live staining

Cerebral organoids (4.5 months old) were cut into 300 μm slices according to the methods described in “Acute slice preparation”. CO slices were perfused at a rate of 2-3 mL/min in oxygenated MitoTracker staining solution containing: ACSF and MitoTracker™ Red CMXRos (Invitrogen™, M7512, 100nM) for 30 minutes at 29–32°C in the dark. After washing three times, samples were fixed in 4% paraformaldehyde overnight at 4°C. Sections of 300μm thickness were embedded into 4% low melting agarose and further cut into 50 μm slices with vibrating-blade microtome (Leica, Germany). Nuclei were counterstained by adding a small drop of ProLong™ Gold Antifade Mountant with DAPI (Invitrogen™, P36935) to each tissue section prior to sealing under glass coverslips. Mountant was left to cure overnight at room temperature and protected from light. Images were captured using Leica TCS SP8 lightning confocal laser scanning microscope (DMI6000) equipped with white light laser (470-670 nm). For imaging of MitoTracker Red CMXRos, 561 nm excitation was used, and emission was acquired at 595 nm by a hybrid detector. Z-stack images were taken at 1 μm intervals and images were collected using LASX software (Lecia Microsystems). For PBMCs and stem cells (iPSCs and H9 hESCs), cells were cultured on Geltrex™-coated 6-well tissue culture plate (Starstedt, 83.3922.005). Cells were incubated in MitoTracker staining solution containing culture media and MitoTracker™ Red CMXRos (Invitrogen™, 100nM) for 30 minutes at 37°C with 5% CO_2_ in the dark. Following incubation, the staining solution was removed, and samples were washed three times (5 minutes per wash) with 1X PBS pH 7.4. After the final wash, FluoroBrite™ DMEM (Gibco™, A1896701) media was added and PBMCs, H9 hESCs and iPSCs were immediately visualized using the EVOS FL Cell Imaging System (Invitrogen™).

### Single-cell dissociation of cerebral organoids

Whole COs were incubated in dissociation solution containing TrypLE Express (Invitrogen™, 12604013) and 0.1 mg/mL of DNase I Solution (STEMCELL Technologies, 07900) for 1 hour at 37°C with 5% CO_2_. During the 1-hour incubation, COs were mechanically triturated (pipetted up and down 5 times) every 15 minutes to dissociate into single cells. At the end of digestion, the digest containing TrypLE Express was neutralized with 5 mL of CO media and the resulting cell solution was filtered twice by passing through cell strainers (Fisherbrand, 22363547, mesh size: 40 μM) attached to a conical tube to further create a single cell suspension. Cells were centrifuged for 1000 RPM for 5 minutes, the supernatant was removed, and the cell pellet was resuspended in 1 mL of CO media for cell count using the Orflo Moxi Flow. The resulting number of cells were used for downstream mitochondrial functional analyses, including ATP and mitochondrial membrane potential measurements. Methods for these experiments are outlined below.

### Mitochondrial membrane potential (MMP) assessment

#### Live imaging for qualitative assessment

Cerebral organoids (4.5 months old) were cut into 100 μm slices according to the methods described in “Acute slice preparation”. Live tissue slices were mounted onto the coverglass bottom surface of 35mm cell imaging dishes (Eppendorf, 145 μm thickness, 0030740009) coated with Cell-Tak Cell and Tissue Adhesive (Corning™, CB40240, 3.5 μg/cm^2^). For PBMCs and stem cells (iPSCs and H9 hESCs), cells were cultured on Geltrex™-coated cover glass of 35mm cell imaging dishes (Eppendorf, 145 μm thickness, 0030740009). Samples were evaluated based on two conditions: basal MMP and inhibited MMP. To visualize basal MMP, samples were incubated in JC-1 staining solution containing culture media and JC-1 Dye (Invitrogen™, Mitochondrial Membrane Potential Probe, T3168, 1 μg/mL) for 30 minutes and 2 drops/mL of NucBlue™ Live ReadyProbes™ Reagent (Invitrogen™) for 20 minutes at 37°C, 5% CO_2_ in the dark. To visualize MMP inhibition, samples were treated with carbonyl cyanide 4-(trifluoromethoxy) phenylhydrazone (FCCP, abcam, ab120081) prepared in culture media at 100 uM concentration for 15 minutes at 37°C, 5% CO_2_ in the dark. After FCCP treatment, samples were incubated in JC-1 staining solution for 30 minutes and 2 drops/mL of NucBlue™ Live ReadyProbes™ Reagent (Invitrogen™, R37605) for 20 minutes at 37°C, 5% CO_2_ in the dark. Following staining, samples were washed three times (5 minutes per wash) with 1X PBS pH 7.4 to remove excess substrate. After the final wash, FluoroBrite™ DMEM (Gibco™) media was added to the imaging dish and samples were immediately visualized using the Leica TCS SP8 lightning confocal laser scanning microscope (DMI6000) equipped with white light laser (470-670 nm) and stage-top humidified incubation system set at 37°C and 5% CO_2_. For imaging of JC-1 dye, 485 ± 11 nm excitation was used, and emission was acquired at 530 ± 15 and ≥590 nm by two hybrid detectors using LASX software (Lecia Microsystems). See Table S4 for biological and technical replicates.

#### Fluorescence spectroscopy for semi-quantitative assessment

Single cells from PBMCs, stem cells (iPSCs and H9 hESCs) and COs were seeded at a density of 200,000 cells per 100 uL FluoroBrite™ DMEM (Gibco™) media per well in a 96-well black polystyrene plate (Greiner CELLSTAR^®^, 655079). Single cells were evaluated based on two conditions: basal MMP (with JC-1 only) and inhibited MMP (100 uM FCCP + JC-1) as described above. Fluorescence readings from FCCP-treated and untreated samples were acquired using a Synergy H1 microplate reader (BioTek^®^ Instruments, Inc., 253147) and Gen5 Software with the following fluorescence excitation/emission settings: red J-aggregates (488 nm/595 nm) and green monomers (488 nm/535 nm). The ratio of fluorescence intensity of red to green is an indicator of the MMP. See Table S4 for biological and technical replicates.

### ATP measurement

ATP levels were measured using CellTiter-Glo Luminescent Cell Viability Assay (Promega, G7570) according to manufacturer’s instructions. Single cells from PBMCs, stem cells (iPSCs and H9 hESCs) and COs were seeded at a density of 50,000 cells per 100 μL culture media per well in a 96-well white polystyrene plate (Greiner CELLSTAR^®^, 655083). An ATP standard curve was generated using ATP disodium salt (Sigma-Aldrich, A7699) ranging from 0 μM to 1 μM with 100 μL of 1μM ATP solution containing 10^−10^ moles ATP. After addition of CellTiter-Glo^®^ Reagent (100 uL) to each well, contents were mixed on an orbital shaker for 2 minutes and luminescence readings from experimental samples and ATP standards were acquired using a Synergy H1 microplate reader (BioTek^®^ Instruments, Inc., 253147) and Gen5 Software. See Table S4 for biological and technical replicates.

### Oligomycin Treatment

To confirm whether the main source of ATP production is from OXPHOS or glycolysis in iPSCs and H9 hESCs, cells were treated with 1 μM of oligomycin for 30 minutes in MEF-conditioned media. The concentration and time have been established in many Seahorse assays to inhibit OXPHOS-linked ATP production by blocking complex V ^27^. Following treatment, cells were immediately seeded at a density of 50,000 cells per 100 μL culture media per well in a 96-well white polystyrene plate (Greiner CELLSTAR^®^, 655083) and ATP production was evaluated using methods as described above.

### Luminex Technology Multiplex Human Oxidative Phosphorylation (OXPHOS) Magnetic Bead Panel

The assembly of mitochondrial complexes I to V in PBMCs, stem cells (H9 hESCs and iPSCs) and COs derived from H9 hESCs and iPSCs were measured using the Human Oxidative Phosphorylation Magnetic Bead Panel Milliplex MAP Kit (Millipore–H0XPSMAG-16K), following the manufacturer’s instructions. Briefly, cell or tissue pellets were resuspended in Cell/Mitochondrial Lysis Buffer (as recommended by Millipore) with protease inhibitors (1:100, EMD Chemicals, Catalog #535140) and phosphatase inhibitors (1:50, EMD Chemicals, Catalog #524629). Protein lysates (500 μg/mL) were incubated with specific proprietary capture antibodies raised against the assembled OXPHOS protein complexes (I, II, III, IV and V) for 2 hours at room temperature. The samples were then incubated with biotinylated secondary detection antibodies for 1 hour at room temperature followed by incubation with streptavidin phycoerythrin conjugate for 30 minutes, which is the reporter molecule to complete the reaction. Samples were ran in five technical replicates. The intensity of the fluorescent signal was acquired using the Luminex Magpix system (Luminex Corporation xMAP Technology) and xPONENT software and analyzed using the MILLIPLEX Analyst 5.1 software. Results were expressed as normalized median fluorescence intensity. See Table S4 for biological and technical replicates.

### Fluorescent immunohisto/cytochemistry for OXPHOS

Cerebral organoid cryosections (14 μm) thickness or cells (PBMCs, H9 hESCs and iPSCs) grown on glass coverslips were used for immunofluorescent staining. For tissues, samples were incubated in permeabilization/blocking solution (1X PBS pH 7.4, 10% Normal Donkey Serum, 2% BSA, 0.5% Triton X-100) for 1 hour at RT in a humidified chamber. For cells, samples were permeabilized with 0.3% Triton X-100 (Sigma-Aldrich) in 1X PBS pH 7.4 for 15 minutes at room temperature and incubated in blocking solution (0.1% Tween-20 in 1X PBS pH 7.4, 1% BSA and 5% Normal Donkey Serum) for 1 hour at room temperature. Proteins of interest were labeled with the following primary antibodies: NDUFS3 (mouse, abcam, ab110246, 1:100), SDHA (mouse, abcam, ab14715, 1:200), UQCRC1 (mouse, abcam, ab110252, 1:50), COXIV (mouse, abcam, ab33985, 1:200), ATP Synthase β (mouse, Invitrogen, A-21351, 1:500) and TOMM20 (rabbit, abcam, ab186734, 1:100) with specific combinations (NDUFS3+TOMM20, SDHA+TOMM20, UQCRC1+TOMM20, COXIV+TOMM20 and ATP Synthase β+TOMM20) diluted in antibody blocking solution (1X PBS pH 7.4, 0.1% Tween-20, 1% BSA, and 5% Normal Donkey Serum) and incubated overnight at 4°C. Following overnight incubation, samples were washed three times (5 minutes per wash) with 0.1% Tween-20 in 1X PBS pH 7.4 and then incubated overnight at 4°C (for tissues) or 1 hour at RT (for cells) with the appropriate secondary antibodies: Donkey anti-Mouse IgG (H+L) Highly Cross-Adsorbed Secondary Antibody Alexa Fluor Plus 488 (Invitrogen, A23766) and Donkey anti-Rabbit IgG (H+L) Highly Cross-Adsorbed Secondary Antibody Alexa Fluor Plus 647 (Invitrogen, A32795) diluted 1:750 in antibody blocking solution. Following secondary antibody incubation, samples were washed three times in 1X PBS pH 7.4 with 0.1% Tween-20. Samples were counterstained with ProLong™ Gold Antifade Mountant with DAPI (Invitrogen™) and left to cure overnight at room temperature protected from light. Images were captured using Leica TCS SP8 lightning confocal laser scanning microscope (DMI6000) equipped with white light laser (470-670 nm) and collected using LASX software (Lecia Microsystems). See Table S4 for biological and technical replicates.

### Transmission Electron Microscopy

Mitochondrial number and morphology for PBMCs, stem cells (H9 hESCs and iPSCs) and COs derived from H9 hESCs and iPSCs were evaluated using Transmission Electron Microscopy. Samples were fixed with primary fixation buffer (0.1M phosphate buffer pH 7.2, 4% paraformaldehyde, 1% Glutaraldehyde) for 2 hours at room temperature and replaced with fresh fixative overnight at 4°C. Following overnight incubation, samples were washed three times for 20 minutes each with 0.1M phosphate buffer at room temperature and underwent a secondary fixation with 1% osmium tetroxide in 0.1M phosphate buffer for 1 hour at room temperature in the dark. The samples were again washed with 0.1M phosphate buffer (pH 7.2) three times for 10 minutes each. Dehydration steps were performed on the samples by using a graded series of ethanol (EtOH) and distilled water at room temperature: 50% EtOH twice for 10 minutes each, 70% EtOH twice for 10 minutes each, 90% EtOH twice for 10 minutes each, and 100% EtOH three times for 15 minutes each. Once samples were dehydrated, they were washed with a transitional solvent, propylene oxide, twice for 15 minutes each. The samples were then infiltrated with epon resin using a graded series of epon and propylene oxide mixture at room temperature: 1) one part Epon resin mixed with two parts 100% propylene oxide for 1 hour using an agitator; 2) two parts Epon resin mixed with one part 100% propylene oxide for 3 hours using an agitator; 3) 100% epon overnight using an agitator and finally; 4) fresh 100% epon resin for 2 hours. Once infiltration was complete, the samples were placed in a Beem embedding capsule and polymerized at 60°C for 48 hours. After complete polymerization, the solid resin block containing the samples were sectioned on a Reichert Ultracut E microtome to 90nm thickness and collected on 200 mesh copper grids. The sections were stained using saturated uranyl acetate for 15 minutes, rinsed in distilled water, followed by Reynold’s lead citrate for 15 minutes and rinsed again in distilled water. The sections were examined and photographed using a FEI Talos L120C transmission electron microscope at an accelerating voltage of 80kV.

## ACKNOWLEDGMENTS

We thank Wendy Horsfall for assistance with mitochondrial DNA haplogroup and sex characterization experiments. A.D. was supported by a Precision Medicine (PRiME) Fellowship, Ontario Graduate Scholarships and a University of Toronto Fellowship. A.C.A. holds a Canada Research Chair Tier 2 in Molecular Pharmacology of Mood Disorders. J.M.B. holds a Canada Research Chair Tier 1 in Molecular Psychiatry.

## Funding

This work was supported by grants from Canadian Institutes of Health Research (CIHR) PJT-148568 to J.M.B, FDN-148455 to L.A., FDN-143252 to J.L.W and #505547 to A.C.A, the Canada First Research Excellence Fund/Medicine by Design to L.A. and J.L.W, the Brain and Behavior Research Foundation (NARSAD Independent Investigator Award) to A.C.A. and Neuroscience Catalyst fund to A.C.A.

## AUTHOR CONTRIBUTIONS

A.D. and A.C.A. designed the study and wrote the manuscript. A.D. performed clinical characterization of the donor, blood collection and processing, iPSC generation from PBMCs, iPSC and cerebral organoid characterization, mitochondrial function experiments, analysis and interpretation of the results. A.E. perform electrophysiology experiments, analyzed the data and summarized the results. A.S. and J.G. in L.A. and J.L.W. laboratories generated and cultured the cerebral organoids and prepared cryosections. J.J.H. stabilized the iPSC line for cerebral organoid generation. N.S. and A.W. performed and evaluated the sequencing of mitochondrial DNA. J.M.B., L.A. and A.C.A. supervised the study. All authors participated in the interpretation of data, in revising the manuscript, and decision to publish the results.

## COMPETING INTEREST

The authors declare no competing interests.

## Notes

### Competing Interest Statement

The authors have declared no competing interest.

